# Diverse incubation rhythms in a facultatively uniparental shorebird – the Northern Lapwing

**DOI:** 10.1101/324426

**Authors:** Martin Sládeček, Eva Vozabulová, Miroslav E. Šálek, Martin Bulla

## Abstract

Incubation by both parents is the most common form of care for eggs. Although the involvement of the two parents may vary dramatically between and within pairs, as well as over the day and breeding season, detailed description of this variation (especially in species with variable male contribution to incubation) is rare. Here, we continuously video-monitored 113 nests of Northern Lapwing *Vanellus vanellus* over the breeding season to reveal the diversity of incubation rhythms and parental involvement. We found great between-nest variation in the overall nest attendance (68 –94%; median = 87%) and in how much males incubated (0 – 37%; median = 13%). Notably, the less the males incubated, the lower was the overall nest attendance, even though females partially compensated for the males’ decrease. Overall, incubation rhythms changed little over the season and incubation period. However, as nights shorten with the progressing breeding season, the female longest night incubation bout shortened too. Essentially, nest attendance was highest, incubation bouts longest, exchange gaps shortest and male involvement lowest during the night. Also, males tended to incubate the most after sunrise and before sunset. To conclude, we revealed strong circadian rhythms and remarkable between nest differences in Northern Lapwing incubation (especially in male involvement), yet despite seasonal environmental trends (e.g. increasing temperature) Lapwing incubation rhythms remained relatively stable over the season and incubation period.

## INTRODUCTION

In majority of avian species both parents incubate the eggs (Deeming 2002). Yet, species vary greatly in how parents divide and time their incubation (Kendeigh 1952; Skutch 1957; Bulla et al. 2016b).

In some species, both sexes share incubation duties nearly equally (Coulson & Wooller 1984; Bulla et al. 2014; Bulla et al. 2016b); in others, one sex dominates the incubation (Afton 1980; Hawkins 1986; Reid et al. 2002; Kleindorfer et al. 2015). In some species, such as seabirds, one parent sits on the nest continuously for several days (e.g., Johnstone & Davis 1990; Weimerskirch 1995; Gauthier-Clerc et al. 2001); others, one parent sits continuously on the nest only for few hours (Grant 1982; Blanken & Nol 1998; Wiebe 2008; Bulla et al. 2014) or even only for a few minutes (Bartlett et al. 2005). Thus, although the general between-species difference in how parents divide and time their incubation is somewhat known, detailed descriptions of how parents divide and time their incubation over the day and season (i.e. as ambient temperatures and predation pressure change) are rare (Bambini et al. in press; Coulson & Wooller 1984; Pedler et al. 2015; Bulla et al. 2016b; Bulla et al. 2017a; Zhang et al. 2017), and these descriptions are mostly limited to species with incubation bouts lasting several days (see above). Moreover, although between- and within-pair differences in incubation rhythms might be considerable (Bulla et al. 2014; Bulla et al. 2016b), and in extreme cases one parent may even desert its incubating partner (Bulla et al. 2017a), detailed description of such between- and within-pair differences is often also lacking.

Here, we used continuous video-monitoring to describe incubation rhythms of the Northern Lapwing *Vanellus vanellus,* a common Palearctic shorebird with variable male contribution to incubation (Liker & Szekely 1999; Grønstøl 2003; Jongbloed et al. 2006). Current knowledge about incubation of Northern Lapwings is mostly based on brief sampling periods of a few hours (Liker & Szekely 1999; Grønstøl 2003; Lislevand et al. 2004; Lislevand & Byrkjedal 2004; but see Jongbloed et al. 2006). Subsequently, we know little about the daily and seasonal variation in how sexes divide their incubation duties between and within pairs. Also, we know little about how male incubation changes over the season.

We specifically investigated (1) between-nest variation in overall nest attendance (proportion of observed time a nest was incubated), (2) how male incubation related to this between-nest variation in nest attendance and (3) how incubation rhythm (female and male contribution) changed within a day, throughout the incubation period and season.

## Methods

### Data collection

In 2015 and 2016 during April and May we monitored incubation of Northern Lapwings in České Budějovice basin, Doudlebia, Czech Republic (49.25°N, 14.08°E), on approximately 40 square kilometres of agricultural landscape. We searched for nests by systematically scanning fields and meadows with telescopes, or by walking through areas with high nest densities. If a nest was found during laying, we estimated its start of incubation by assuming that females laid one egg per day and started incubation when the clutch was complete (usually four, rarely three eggs). If a nest was found with a full clutch, we estimated its start of incubation based on the median height and angle at which the eggs floated in water and assuming 27 days long incubation period (Van Päässen et al. 1984).

We monitored incubation with a custom designed video recording system (Jan Petru, Czech Republic), consisting of an external lens (Ø 2 cm, length 4 cm) mounted on a ^~^30 cm long twig and placed 1.5 meters from the nest in a southward direction to minimize the time the lens faced the sun, which would have overexposed the videos and made individuals hard to recognize. The digital recorder stored videos in 10-15 frames per second in 640 × 480 pixels resolution for about four days. The system was powered by a 12-V, 44-Ah battery buried together with the recorder (in a waterproof case) under the ground.

### Extraction of incubation behaviour

We extracted incubation behaviour from video recordings in AVS Media Player (http://www.avs4you.com/AVS-Media-Player.aspx) by noting date and time (to the nearest second) when a bird came to the nest (both legs in the nest) or left the nest. We thus define incubation as both, sitting on the eggs (warming) or standing above them (turning them or shading them from direct sunlight). We distinguished females and males via individual and sex-specific plumage traits such as crest length, extent of melanin ornaments on the face and breast (Meissner et al. 2013). We further noted any disturbance caused by the field team, agricultural work, general public or interaction with other animals (note that only bouts with disturbance from the field team were excluded from the analyses). Bouts with technical difficulties and with low visibility, when parents were hard to recognize (e.g. during direct sunlight or heavy rains), were classified as uncertain and excluded from the analyses (see Supplementary Actograms for details, raw incubation data and extracted incubation bouts; Sládeček & Bulla 2018).

### Definition of incubation variables

We define nest attendance as the proportion of time when a nest was actually incubated. Specifically, ‘overall nest attendance’ indicates nest attendance for the whole time a nest was monitored; ‘daily nest attendance’ indicates attendance for a particular day and nest; ‘hourly nest attendance’ indicates attendance for a particular hour in a particular day and nest. Female or male nest attendance denotes proportion of incubation by a particular sex during a respective time interval (e.g. overall, day, hour or incubation bout).

Furthermore, we define incubation bouts as the total time allocated to a single parent (that is, the time between the arrival of a parent at the nest and its departure, followed by the incubation of its partner) and exchange gaps as the time between the departure of one parent from the nest and the return of its partner.

Last, responsibility indicates sum of all recorded incubation bouts for a given parent and nest.

### Sample sizes

We have monitored 107 nests (46 in 2015 and 61 in 2016) for median 3 days (range: 1-7 days); in addition, we included another 6 nests from a different study with median 14 days (range: 8-22 days Bulla et al. 2016a; Bulla et al. 2016b). However, not all incubation data and nests were suitable for all analyses. For the analyses of overall nest attendance, we used only nests with at least two complete days of recorded incubation (*N* = 60 nests with median of 3 days per nest; range: 2-20 days). For the analyses of daily nest attendance, we used only nests with at least one day of recorded incubation (*N* = 191 days from 78 nests with median of 2 days per nest; range: 1-20 days). For both, nest level and daily nest attendance data, we used only days monitored for more than 90% of day. For the analyses of hourly nest attendance, we used only nests with at least 24 hours of recording and only hours with complete incubation recording (*N* = 113 nest with median of 61 hours, range: 24-482 hours). We used the same nests (but excluding uniparental ones) for the analyses of incubation bouts and exchange gaps (*N* = 107 nest with median of 20 incubation bouts and exchange gaps per nest, range: 1-297 bouts and exchange gaps).

### Statistical analysis

All procedures were performed in R version 3.3.0 (R-Core-Team 2017). General linear models were fitted using ‘lm’ function (R-Core-Team 2017) and mixed effect models using ‘lmer’ functions from the ‘lme4’ R package (Bates et al. 2015). For each model parameter we report effect size and model predictions as medians and the Baysian 95% credible intervals (95%CI) represented by 2.5 and 97.5 percentiles from the posterior distribution of 5 000 simulated or predicted values obtained by the ‘sim’ function from the ‘arm’ R package (Gelman et al. 2016). We estimated the repeatability of female and male nest attendance using the ‘rpt’ function from the ‘rptR’ R package (Nakagawa & Schielzeth 2010), restricted maximum likelihood method (REML), gaussian model, and 5 000 bootstrapped runs (Nakagawa & Schielzeth 2010). The specification of each model are described in detail in the Supplementary Information (Sládeček & Bulla 2018).

## RESULTS

### Overall nest attendance

Northern Lapwing parents incubated their eggs 87% of time (median, range: 68 – 94%, *N* = 60 nests with more than 2 days of recording; Figure 1a, dark red and blue). Actual incubation bouts covered 98% of observed time (median, range: 95 – 100%, *N* = 55 nests incubated by both parents; Figure 1a, red and blue). In other words, one parent was nearly always responsible for the nest (sum of incubation bouts). Exchange gaps thus accounted for only 2% of observed time (median, range: 0.3 – 5%, *N* = 55 nests incubated by both parents).

**Figure. 1.**
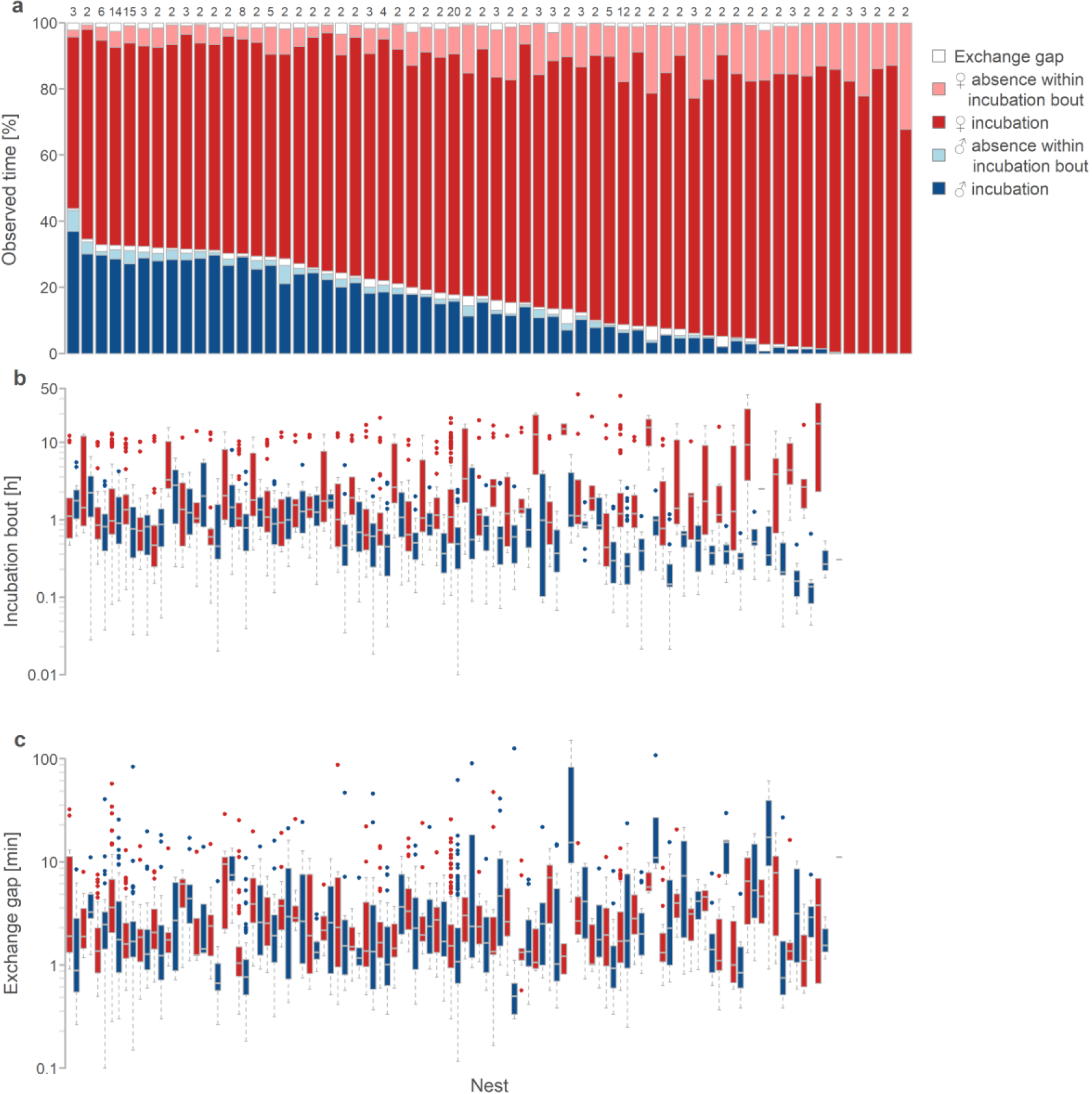
Between and within nests variation in incubation. **a**, Between- and within-nest variation in nest attendance. Each bar represents one nest and proportion of female incubation bouts (red), male incubation bouts (blue) and exchange gaps (gaps preceding female incubation bouts are above female bars and those preceding male incubation bouts are above male bars). Dark colours indicate actual incubation (individual sitting on the nest) and light colours indicate absence of a parent (no incubation) within its incubation bouts. Numbers above the bars indicate number of days with incubation data. Nests (bars) are ordered from the highest to the lowest male nest-attendance (*N* = 60 nests). **b, c**, Between- and within-nest variation in incubation bouts (**b**) and exchange gaps (**c**) according to sex (female in red, male in blue). Each pair of box plots (female and male) corresponds to the nest (bar) in **a** (*N* = 2239 bouts and gaps from 55 biparental nests). Box plots depict median (vertical line inside the box), 25-75th percentiles (box), 25th and 75th percentiles minus or plus 1.5× interquartile range, respectively, or the minimum and maximum value, whichever is smaller (whiskers) and outliers (circles).

Females dominated the incubation, incubating the eggs 72% of observed time (median, range: 52 – 87%, *N* = 60 nests with more than 2 days of recording; Figure 1a in dark red) and being responsible for the nest, that is their incubation bouts covered, 82% of observed time (median, range: 54 – 100%; Figure 1a in dark and light red; note that 100% represents 5 nests incubated solely by females). Females were absent from the nest during their incubation bouts 10% of time (median, range: 2 – 32%; Figure 1a in light red). In contrast, males attended the nests 13% of time (median, range: 0 – 37%; Figure 1a in dark blue) and were responsible for the nest 15% of time (median, range: 0 – 43%; Figure 1a in dark and light blue). In 55 nests with male incubation, males were absent from the nest during their incubation bouts 7% of time (median, range: 0 – 22%; Figure 1a in light blue). Note that overall female nest attendance was always higher than that of males and the two strongly negatively correlated (*r* = - 0.87; Figure S3a).

Overall nest attendance decreased as male nest attendance decreased (Figure 1a and Figure 2a, Table S1). Nevertheless, females ‘partially compensated’ for this decrease. As female responsibility for the nest increased, their nest attendance increased as well, but less then would be expected under the ‘full compensation’ (Figure 1a and 2b, Table S2; note that in Figure 2b hypothetical full compensation is indicated by dashed line and in case of no compensation the points would be parallel to x-axis).

**Figure. 2.**
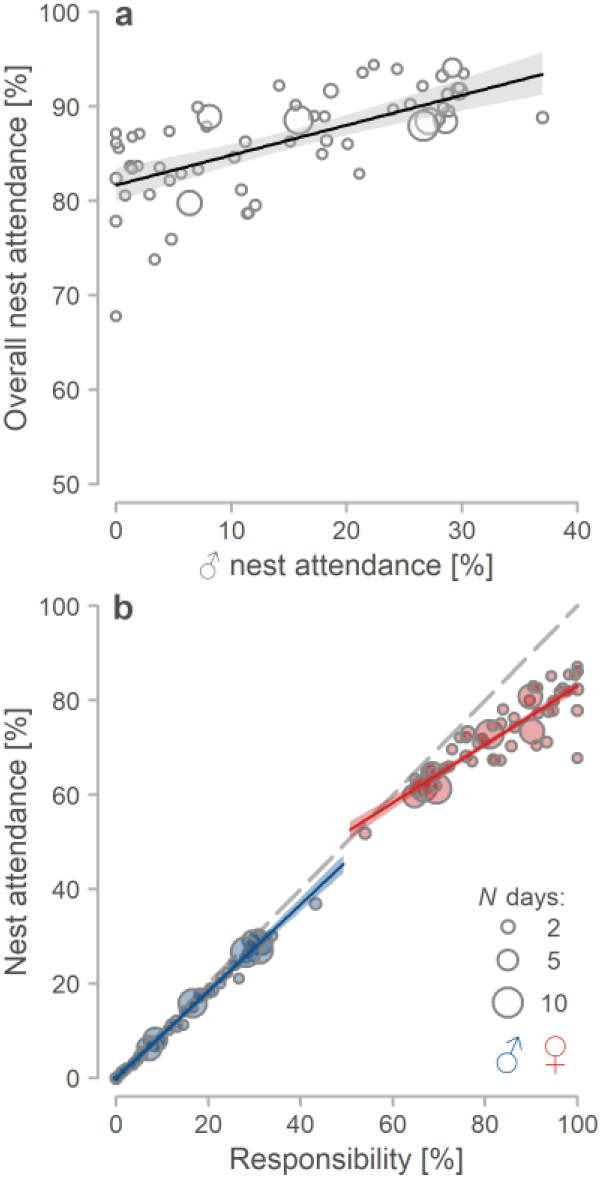
Contribution of female and male to incubation. **a**. Relationship between male nest attendance and overall nest attendance (*N* = 60 nests). **b**. Relationship between responsibility (proportion of all parent’s incubation bouts within observed time) and nest attendance for females (red) and males (blue; *N* = 120 parents from 55 biparental nests). **a, b**, Circles represent individual nests (**a**) or parents (**b**) and their size number of days with incubation data. Lines with shaded areas indicate model prediction with 95% credible intervals based on the joint posterior distribution of 5,000 simulated values based on model outputs (Table S1 and S2) and generated by the ‘sim’ function in R (Gelman et al. 2016). Included are only nests with at least two days of incubation data and days with at least 90% of recording. Dashed line in **b** indicates full compensation for the reduced care of a partner.

### Daily nest attendance

Daily nest attendance mirrored the overall nest attendance (median = 88%, range: 50 – 98%, *N* = 191 days from 78 nests) and also increased with male nest attendance (Figure S1, Table S3). Daily nest attendance was repeatable in females (0.54, 95%CI: 0.37 – 0.67), as well as in males (0.7, 95%CI: 0.58 – 0.8). Although daily nest attendance was unrelated to the day in incubation period or day when the nest started within the breeding season (Table S3), it varied strongly within a day, being highest and nearly instantaneous during the night and lowest during the day (Figure 3a - in dark grey). Females were nearly sole incubators at night, while assisted by males during the day (Figure 3a and 3b; Table S3). Specifically, female incubation dropped (male incubation peaked) after sunrise and in males peaked also before sunset (Figure 3a, b; Table S3).

**Figure. 3.**
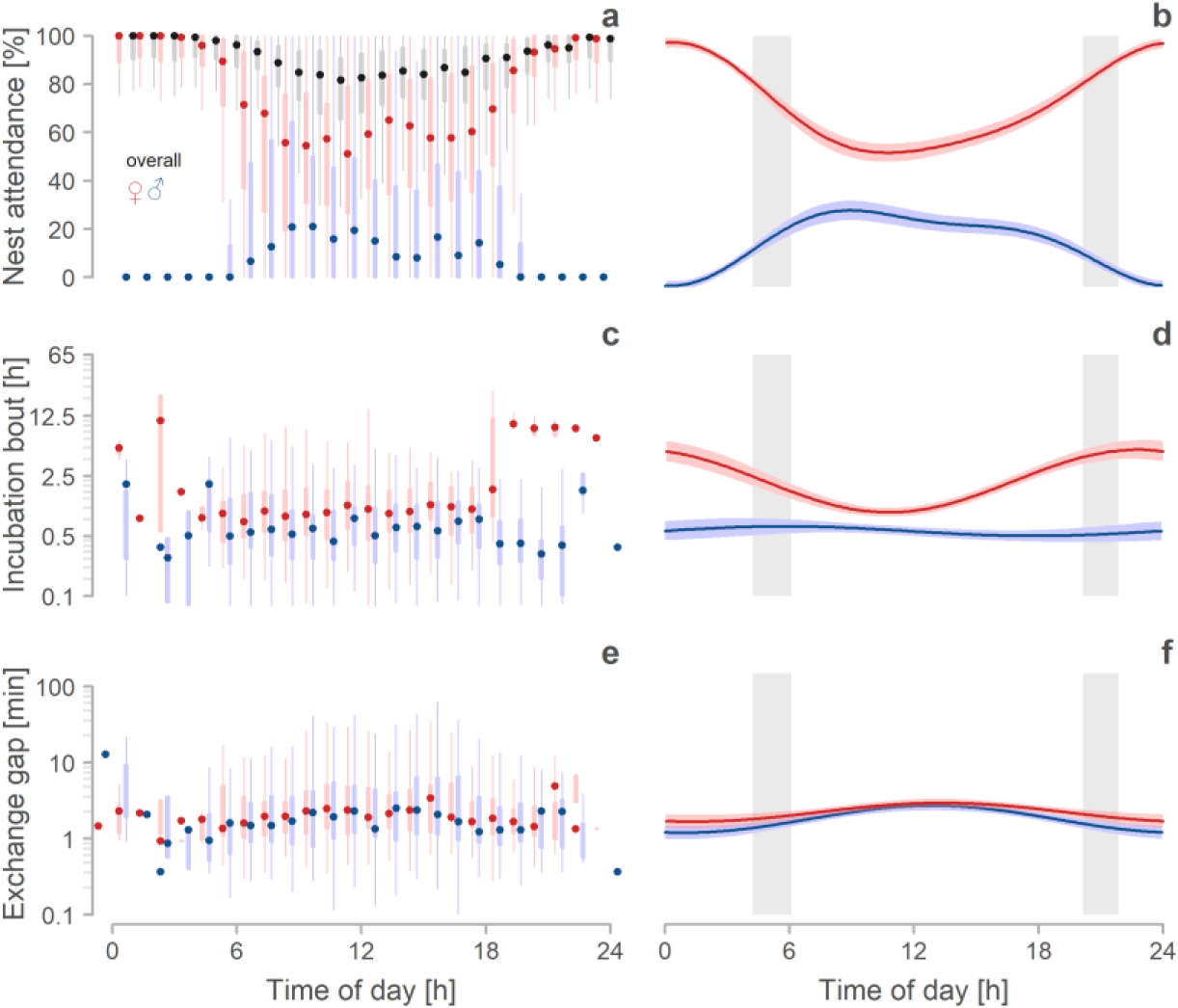
Daily changes in incubation behaviour. **a, c, e,** Daily variation in overall (dark grey), female (red) and male (blue) nest attendance (**a**), bout length (**c**) and exchange gap (**e**). Points depict hourly median weighted by sample size for each nest. Thicker lines indicate 25–75th percentiles, thinner lines 25th and 75th percentiles minus or plus 1.5× interquartile range, respectively, or the minimum and maximum value, whichever is smaller. Note that outliers are not depicted. Included are only hours with complete incubation record (*N* = 7933 hours from 113 nests; median 61 hours per nest, range: 24-482) and complete incubation bouts (*N* = 3184 bouts from 107 biparentally incubated nests; median 20 bouts and exchange gaps per nest, range: 1-297). **b, d, f,** Predicted relationships between time of day and nest attendance (**b**), bout length (**d**) and exchange gap length (**f**) according to sex (female in red, male in blue). Lines with shaded areas indicate model prediction with 95% credible intervals based on the joint posterior distribution of 5,000 simulated values from mode outputs (Table S4, S5 and S8) and generated by the ′sim′ function in R (Gelman et al. 2016). Grey bars indicate the period between the earliest and the latest sunrise and sunset during incubation monitoring.

### Incubation bouts and exchange gaps

Incubation bouts of biparental nests lasted 44 min (median, range: 1s – 42h, *N* = 3184 bouts from 107 nests) and varied greatly between and within nests (Figure 1b, Supplementary Actograms) and especially over the day (Figure 3c, d); bouts (especially of females) were longer during the night than during the day (Figure 3c, d). Female incubation bouts lasted 1h (median, range: 1min – 42h; N = 1518) whereas male incubation bouts lasted 32min (median, range: 1s – 7.9h; N = 1666). Notably, on average and regardless of time, female bouts were always longer than those of males (Figure 3c, d; Table S5), although during the daytime females had shorter median incubation bout than males in 30% (Figure S3b; also note that female and male bout were uncorrelated). Overall, incubation bouts were similar across incubation period and unrelated to the day when the nest started within breeding season, but note the tendency in males for shorter incubation bouts as incubation period progressed (Figure S2, Table S5). Essentially, as breeding season progressed and nights became shorter, female night incubation bouts shortened too (Figure 4a, Table S6). Also, median bout length of females and males positively correlated with their nest attendance (Figure 1 and Figure 4b; Table S7).

**Figure. 4.**
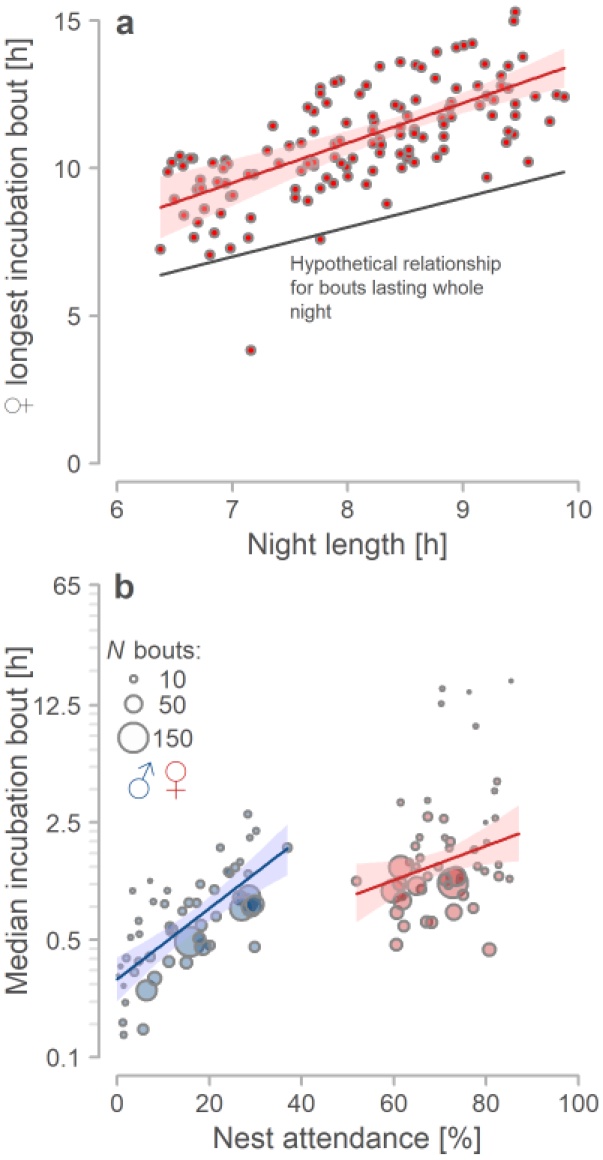
Bout length correlates. **a,** Longest female incubation bout during particular night in relation to the night length - defined as time when sun is > 6° under the horizon. Points indicate bouts (*N* = 133 bouts from 55 nests; median 2 nights per nest, range 1-14 night; included are only bouts with at least 60% of their length in night). Thick grey line indicates a situation when incubation bouts would last precisely the whole night. **b,** Median bout length in relation to sex (females in red, males in blue) and nest attendance. Circles represent individual parents, their size number of days with incubation data. (*N* = 110 parents from 55 biparentally incubated nests with more than two days of continuous recording). **a, b**, Lines with shaded areas indicate model prediction with 95% credible intervals based on the joint posterior distribution of 5,000 simulated based on model outputs (Table S6 and S7) and values generated by the ‘sim’ function in R (Gelman et al. 2016).

Exchange gaps lasted 1.9 min (median; range 6s – 2.5h; *N* = 3184 exchange gaps from 107 nests). Length of exchange gaps was unrelated to the day in incubation period and the day when the nest started within the season. However, exchange gaps fluctuated over the day, being longest in the middle of the day (noon median = 2.49 min, range: 0.13-39.5 min, interquartile range: 1.32-5.32), and shortest during night, specifically during mornings (5:00 o′clock median = 1.16 min, range: 0.3519.3 min, interquartile range: 0.68-2.05) and evenings (19:00 o′clock median = 1.41 min, range: 0.2887.5, interquartile range: 0.61-2.63; Figure 3e, f; note that there is only negligible number of exchange gaps during the night). Also, exchange gaps that occurred before female incubation bouts were longer than those before male incubation bouts (Figure 3e, f; Table S8).

## DISCUSSION

Using continuous video monitoring, we quantitatively described incubation rhythms in a Central European population of a common Palearctic shorebird, the Northern Lapwing. We revealed three main aspects of Northern Lapwing incubation rhythms, which we discuss below – (1) nests vary remarkably in overall nest attendance; (2) male contribution to incubation shapes the incubations rhythm, as females are unable to fully compensate for the absence of male care; and (3) the incubation rhythms varied little over the incubation period and season, but varied strongly over the day.

### Overall incubation rhythm

First, overall nest attendance was 87% (median; range: 67 to 94%; Figure 1a), which is in line with findings from other Lapwing populations (Liker & Szekely 1999; Grønstøl 2003; Lislevand & Byrkjedal 2004; Jongbloed et al. 2006). Yet, it remains unclear why Northern Lapwings do not incubate more, given that closely related species can achieve higher nest attendance even when incubating uniparentally (Løfaldli 1985; Kålås 1986). Perhaps breeding in a temperate climate does not require continuous incubation attendance (but see temperate species in Bulla et al. 2016b). Essentially, the between nest variability in nest attendance – which can be as much as a 6.5 hours difference per day – seems huge and is much larger than nest attendance fluctuations known to influence embryo development (Hepp et al. 2006; Carter et al. 2014; Bueno-Enciso et al. 2017), or length of the incubation period (Grønstøl 2003; Martin et al. 2007; Carter et al. 2014; Bueno-Enciso et al. 2017).

Moreover, we found that nest attendance positively correlated with length of incubation bouts (Figure 4b) and that exchange gaps between incubation bouts were short (median = 1.9min; Figure 3c). These findings suggest that the departures of a parent from its nest (within its incubation bout) do not trigger their often nearby partner to come and incubate (Parish & Coulson 1998; Lislevand et al. 2004). Hence, in Northern Lapwings other (to date unknown) cues drive the decision of parents to return to the nest and incubate. These findings resemble those from other species (Johnstone & Davis 1990; Weimerskirch 1995; Bulla et al. 2014), albeit that in most of those the off-nest partner is far from the nest.

### Male contribution to incubation

Second, we found immense variation in male contribution to incubation. Male nest attendance varied from 0 to 37% of observed time, with median of only 13% (Figure 1). Such male contribution is lower than in other Northern Lapwing populations (27 % in Liker & Szekely 1999; 21 % in Lislevand 2001; 19 % in Lislevand & Byrkjedal 2004; 19 % in Jongbloed et al. 2006). This difference may result from the daytime only monitoring of incubation in most previous studies, since if we include only day-light period, male nest attendance in our data rise to 18% (median).

The low male nest attendance across Northern Lapwing populations indicate that males invest into parental care differently than by incubating, e.g. by guarding the nest (and the female) against predators (Liker & Szekely 1999; Kis et al. 2000). Importantly, the variation in male nest attendance may reflect the male′s mating status, as some Northern Lapwing males tend to have more than one female (reviewed in: Šálek 2005). We have not recorded mating status, but note that the current evidence for relationship between male mating status and its incubation effort is equivocal and based on non-continuous monitoring (Liker & Szekely 1999; Grønstøl 2003; Lislevand et al. 2004).

Notably, we also found that when male nest attendance was low, female nest attendance was higher, but not enough to fully compensate for the male decrease; hence, with decreasing male attendance, overall as well as daily nest attendance decreased as well (see Figure 2 for overall effects, Figure S1 for daily effects). These findings suggest that, unlike in other species such as geese (Jónsson et al. 2007) with uniparental female incubation and close to 100% nest attendance, the Northern Lapwing females have a limited capacity to incubate continuously. Nevertheless, such limited capacity is in line with nest attendance of other uniparentally incubating shorebirds (Løfaldli 1985; Bulla et al. 2017. Moreover, our finding of partial compensation is in line with previous empirical and theoretical work (McNamara et al. 1999; McNamara et al. 2003; Houston et al. 2005; Harrison et al. 2009). Also note that such partial compensation is feasible only in environments (like in this temperate Northern Lapwing population) where decrease in parental care does not necessarily translate into breeding failure (e.g. due to cooling or overheating of eggs) (Jones et al. 2002; Bulla et al. 2017b).

### Variation in incubation rhythms

Third, incubation rhythms were generally stable during the incubation period and season (Tables S3, S5, S8), but varied strongly within a day (Figure 3). Overall and female nest attendance were highest during the night and lowest during the day; in contrast, males rarely incubated at night and their nest attendance peaked (Figure 3b) after sunrise and before sunset. Similar to the nest attendance, female incubation bouts were long during the night and short during the day, while male incubation bouts were similar across the day (Figure 3b).

The general lack of variation in Northern Lapwing incubation across incubation period and season contrasts with findings from other species where, for example, incubation bouts lengthen over the incubation period and then shorten just before hatching (Bulla et al. 2014; Pedler et al. 2015; Zhang et al. 2017). Note that the lack of variation across incubation period in our study may also reflect lack of statistical power, that is 5-8 days of incubation data at the start and end of incubation period may still not be enough, in face of immense within- and between-nest variation. However, we also found that female night bouts shortened as nights shorten with the progressing breeding season. We propose that this seasonal pattern results from daily incubation rhythm of Lapwings where females take nearly sole responsibility for their nest and incubate continuously with one or few long incubation bouts over the whole night. As nights become shorter, the night incubation bouts also shorten.

So why do Lapwings incubate so differently during the day and during the night? Anti-predation strategy does explain variation in incubation rhythms of shorebirds on a comparative scale; species that rely on camouflage when incubating (i.e. are cryptic) have longer incubation bouts than those which do not (Bulla et al. 2016b). Thus, while Lapwings actively attack predators and have short incubation bouts during the day (Liker & Szekely 1999; Kis et al. 2000), they may rely on crypsis (minimize number of changeovers on the nest) during the night when mammalian predators are active, leading to long incubation bouts. The poor visibility during the night may prohibit Lapwings from attacking mammalian predators. Perhaps, more importantly, attacking a mammal may not deter it from further searching for the eggs. Indeed, in our population all video-recorded egg predation events occurred during the night and by mammals (unpublished data).

Female night incubation is reported in related plover species of genus *Pluvialis* (Bulla et al. 2016a; Bulla et al. 2016b; but see: Cardilini et al. 2015). The lack of male night incubation (Figure 3a) corroborates with findings from Dutch Lapwing populations (Lislevand et al. 2004; Jongbloed et al. 2006). However, we found some night incubation only in 5 males out of 55 incubating males (9%); whereas, in the Dutch population 50% of incubating males incubated at night (Jongbloed et al. 2006). Still, why do males incubate so rarely during the night? One hypothesis suggests that the brighter parent should incubate at night (Ekanayake et al. 2015). However, in Northern Lapwings male is the more ornamented one (von Blotzheim 1999; Meissner et al. 2013) and hence should incubate during the night, but this is not the case.

Notably, the continuous data on 113 nests allowed us to depict distribution of male attendance across day with peaks after sunrise and before sunset (Figure 3a). We speculate that by incubating after sunrise and before sunset males may allow females to replenish their energy stores after and before long night incubation bouts. We thus propose testing whether females lacking male assistance will weigh less and incubate less in the morning, at the end of the day or at the end of the incubation period than females with male contribution to incubation.

### Conclusion

To conclude, with continuous monitoring of Northern Lapwing nests we demonstrate how male contribution to incubation shapes remarkable variability in Lapwing incubation rhythms. We further reveal strong daily fluctuations in the incubation rhythms that further translate to changes in night incubation over the breeding season. We thus not only clarify and specify the current knowledge about Lapwing incubation, but also demonstrate how the advancements in continuous monitoring allow for detailed descriptions of variation in behaviour over multiple time scales (day, incubation period, season), providing thorough insight into the rhythms and tactics of avian reproduction.

### Open data, codes and materials

All available from Open Science Framework http://osf.io/v4vpe (Sládeček and Bulla 2018).

### Authors′ contributions

M.S., E.V. and M.Š. collected the data; M.S. and E.V. extracted incubation from videos, M.S., M.B. and M.Š. conceived the paper, M.S. and M.B. analysed and visualised the data, drafted the paper and with input from M.Š. and E.V. wrote the final paper.

## Acknowledgements

We thank all that helped us in the field, especially to Radka Piálková, Hana Vitnerová, Vojtěch Kubelka, Jana Hronková, Soňa Novotná, Tereza Kejzlarová, and Zuzana Karlíková.

We are also grateful to Kristýna Nohejlová for help with incubation extraction. MB thanks Bart Kempenaers for office space and support and Borka and Majka for patience.

## REFERENCES

Afton, A.D. 1980. Factors Affecting Incubation Rhythms of Northern Shovelers. Condor 82: 132–137.

Bambini, G., Schlicht, E., & Kempenaers, B. Patterns of female nest attendance and male feeding throughout the incubation period in Blue Tits Cyanistes caeruleus. Ibis. doi:10.1111/ibi.12614

Bartlett, T.L., Mock, D.W., & Schwagmeyer, P.L. 2005. Division of Labor: Incubation and Biparental Care in House Sparrows (Passer domesticus). Auk 122: 835–842.

Bates, D., Maechler, M., Bolker, B., Walker, S., Christensen, R.H.B., Singmann, H., Dai, B., & Grothendieck, G. 2015. Fitting Linear Mixed-Effects Models Using lme4. J. Stat. Softw. 67: 1–48.

Blanken, M.S., & Nol, E. 1998. Factors Afecting Parental Behavior in Semipalmated Plovers. Auk 115: 166–174.

von Blotzheim, G. 1999. Handbuch der Vögel Mitteleuropas Band 6: Charadriiformes (1. Teil) (G. von Blotzheim, Ed.). AULA-Verlag, Weisbaden.

Bueno-Enciso, J., Barrientos, R., Ferrer, E.S., & Sanz, J.J. 2017. Do extended incubation recesses carry fitness costs in two cavity-nesting birds? J. F. Ornithol. 88: 146–155.

Bulla, M., Valcu, M., Dokter, A.M., Dondua, A.G., Kosztolányi, A., Rutten, A.L., Helm, B., Sandercock, B.K., Casler, B., Ens, B.J., Spiegel, C.S., Hassell, C.J., Küpper, C., Minton, C., Burgas, D., Lank, D.B., Payer, D.C., Loktionov, E.Y., Nol, E., Kwon, E., & Smith, F. 2016a. Supporting Information for ″Unexpected diversity in socially synchronized rhythms of shorebirds.″

Bulla, M., Valcu, M., Dokter, A.M., Dondua, A.G., Kosztolányi, A., Rütten, A.L., Helm, B., Sandercock, B.K., Casler, B., Ens, B.J., Spiegel, C.S., Hassell, C.J., Küpper, C., Minton, C., Burgas, D., Lank, D.B., Payer, D.C., Loktionov, E.Y., Nol, E., Kwon, E., & Smith, F. 2016b. Unexpected diversity in socially synchronized rhythms of shorebirds. Nature 1: 1–17.

Bulla, M., Valcu, M., Prüter, H., Vitnerová, H., Tijsen, W., Sládeček, M., Alves, J.A., Gilg, O., & Kempenaers, B. 2017a. Flexible parental care: Uniparental incubation in biparentally incubating shorebirds. Scientific Reports

Bulla, M., Valcu, M., Prüter, H., Vitnerová, H., Tijsen, W., Sládeček, M., Alves, J.A., Gilg, O., & Kempenaers, B. 2017b. Flexible parental care: Uniparental incubation in biparentally incubating shorebirds. Scientific Reports. doi:http://dx.doi.org/10.1101/117028

Bulla, M., Valcu, M., Rutten, A.L., & Kempenaers, B. 2014. Biparental incubation patterns in a high-Arctic breeding shorebird: how do pairs divide their duties? Behav. Ecol. 25: 152–164.

Cardilini, A.P.A., Weston, M.A., Dann, P., & Sherman, C.D.H. 2015. Sharing the Load : Role Equity in the Incubation of a Monomorphic Shorebird, the Masked Lapwing ( Vanellus miles). Wilson J. Ornithol. 127: 730–733.

Carter, A.W., Hopkins, W.A., Moore, I.T., & Durant, S.E. 2014. Influence of incubation recess patterns on incubation period and hatchling traits in wood ducks Aix sponsa. J. Avian Biol. 45: 273–279.

Coulson, J.C., & Wooller, R.D. 1984. Incubation under natural conditions in the kittiwake gull, Rissa tridactyla. Anim. Behav. 32: 1204–1215.

Deeming, C. 2002. Avian incubation: behaviour, environment and evolution. Oxford University Press.

Ekanayake, K.B., Weston, M.A., Nimmo, D.G., Maguire, G.S.s, Endler, J.A., & Ku, C. 2015. The bright incubate at night: sexual dichromatism and adaptive incubation division in an open-nesting shorebird.

Gauthier-Clerc, M., Le Maho, Y., Gendner, J.P., Durant, J., & Handrich, Y. 2001. State-dependent decisions in long-term fasting king penguins, Aptenodytes patagonicus, during courtship and incubation. Anim. Behav. 62: 661–669.

Gelman, A., Su, Y.-S., Yajima, M., Hill, J., Pittau, M., Kerman, J. ouni, Zheng, T., & Vincent, D. 2016. Data Analysis Using Regression and Multilevel/Hierarchical Models. CRAN Repository 1–53.

Grant, G.S. 1982. Avian Incubation : Egg Temperature, Nest Humidity, and behavioral thermoregulation in a hot environment. Ornithol. Monogr. 30: 1–82.

Grønstøl, G.B. 2003. Mate-sharing costs in polygynous Northern Lapwings Vanellus vanellus. Ibis (Lond. 1859). 145: 203–211.

Harrison, F., Barta, Z., Cuthill, I., & Székely, T. 2009. How is sexual conflict over parental care resolved? A meta-analysis. J. Evol. Biol. 22: 1800–1812.

Hawkins, L.L. 1986. Nesting behavior of male and female Whistling Swans and implications of male incubation. Wildfowl 37: 5–27.

Hepp, G.R., Kennamer, R.A., & Johnson, M.H. 2006. Maternal effects in Wood Ducks: Incubation temperature influences incubation period and neonate phenotype. Funct. Ecol. 20: 307–314.

Houston, A.I., Székely, T., & Mcnamara, J.M. 2005. Conflict between parents over care. Evolution (N. Y). 20:.

Johnstone, R.M., & Davis, Ll.S. 1990. Incubation routines and foraging-trip regulation in the Grey-faced Petrel Pterodroma macroptera gouldi. Ibis (Lond. 1859). 132: 14–20.

Jones, K., Ruxton, G., & Monaghan, P. 2002. Model parents: is full compensation for reduced partner nest attendance compatible with stable biparental care? Behav. Ecol. 13: 838–843.

Jongbloed, F., Schekkerman, H., & Teunissen, W. 2006. Verdeling van de broedinspanning bij Kieviten. Limosa 79: 63–70.

Jónsson, J.E., Afton, A.D., & Alisauskas, R.T. 2007. Does body size influence nest attendance? A comparison of Ross′s geese (Chen rossii) and the larger, sympatric lesser snow geese (C. caerulescens caerulescens). J. Ornithol. 148: 549–555.

Kålås, J.A. 1986. Incubation schedules in different parental systems in the Dotterel Charadrius morinellus. Ardea 74: 185–190.

Kendeigh, S.C. 1952. Parental care and its evolution in birds.

Kis, J., Liker, A., & Székely, T. 2000. Nest defence by Lapwings: Observation on natural behaviour and an experiment. Ardea 88: 155–164.

Kleindorfer, S., Fessl, B., & Hoi, H. 2015. More Is Not Always Better: Male Incubation in Two Acrocephalus Warblers. Behaviour 132: 607–625.

Liker, A., & Szekely, T. 1999. Parental behaviour in the Lapwing Vanellus vanellus. Ibis (Lond. 1859). 141: 608–614.

Lislevand, T. 2001. Male incubation in Northern Lapwings : effects on egg temperature and potential benefits to females. Ornis Fenn. 78: 23–29.

Lislevand, T., & Byrkjedal, I. 2004. Incubation behaviour in male Northern Lapwing Vanellus vanellus in relation to mating opportunities and female body condition. Ardea 92: 19–30.

Lislevand, T., Byrkjedal, I., Grønstøl, G.B., Hafsmo, J.E., Kallestad, G.R., & Larsen, V.A. 2004. Incubation Behaviour in Northern Lapwings : Nocturnal Nest Attentiveness and Possible Importance of Individual Breeding Quality. Ethology 110: 177–192.

Løfaldli, L. 1985. Incubation Rhythm in the Great Snipe Gallinago media. Holarct. Ecol. 8: 107–112.

Martin, T.E., Auer, S.K., Bassar, R.D., Niklison, A.M., & Lloyd, P. 2007. Geographic variation in avian incubation periods and parental influences on embryonic temperature. Evolution (N. Y). 61: 2558–2569.

McNamara, J.M., Gasson, C.E., & Houston, A.I. 1999. Incorporating rules for responding into evolutionary games. Nature 401: 368–371.

McNamara, J.M., Houston, A.I., Barta, Z., & Osorno, J.L. 2003. Should young ever be better off with one parent than with two?. Behav. Ecol. 14: 301–310.

Meissner, W., Wójcik, C., Pinchuk, P., & Karlionova, N. 2013. Part 9 : Ageing and sexing the Northern Lapwing Vanellus vanellus. Wader Study Gr. Bull. 120: 32–36.

Nakagawa, S., & Schielzeth, H. 2010. Repeatability for Gaussian and non-Gaussian data: a practical guide for biologists. Biol. Rev. Camb. Philos. Soc. 85: 935–56.

Van Päässen, A.G., Veldman, D.H., & Beintema, A.J. 1984. A simple device for determination of incubation stages in eggs. Wildfowl 35: 173–178.

Parish, D.M.B., & Coulson, J.C. 1998. Parental investment, reproductive success and polygyny in the lapwing, Vanellus vanellus. Anim. Behav. 56: 1161–1167.

Pedler, R.D., Weston, M.A., & Bennett, A.T.D. 2015. Long incubation bouts and biparental incubation in the nomadic Banded Stilt. Emu 116: 75–80.

R-Core-Team. 2017. R: A Language and Environment for Statistical Computing.

Reid, J.M., Monaghan, P., & Ruxton, G.D. 2002. Males matter: The occurrence and consequences of male incubation in starlings (Sturnus vulgaris). Behav. Ecol. Sociobiol. 51: 255–261.

Skutch, A.F. 1957. the Incubation Patterns of Birds. Ibis (Lond. 1859). 99: 69–93.

Sládeček, M., & Bulla, M. 2018. Supporting information for “Diverse incubation rhythm in a facultatively uniparental shorebird - the Northern Lapwing.”

Šálek, M. 2005. Polygamous breeding of Northern Lapwings (Vanellus vanellus) in southern Bohemia, Czech Republic. Sylvia 41: 72–82.

Weimerskirch, H. 1995. Regulation of foraging trips and incubation routine in male and female wandering albatrosses. Oecologia 102: 37–43.

Wiebe, K.L. 2008. Division of labour during incubation in a woodpecker Colaptes auratus with reversed sex roles and facultative polyandry. Ibis (Lond. 1859). 150: 115–124.

Zhang, L., Shu, M., An, B., Zhao, C., Suo, Y., & Yang, X. 2017. Biparental incubation pattern of the Black-necked Crane on an alpine plateau. J. Ornithol. 158: 697–705.

